# Memory effects and static disorder reduce information in single-molecule signals

**DOI:** 10.1101/2022.01.13.476256

**Authors:** Kevin Song, Dmitrii E. Makarov, Etienne Vouga

## Abstract

A key theoretical challenge posed by single-molecule studies is the inverse problem of deducing the underlying molecular dynamics from the time evolution of low-dimensional experimental observables. Toward this goal, a variety of low-dimensional models have been proposed as descriptions of single-molecule signals, including random walks with or without conformational memory and/or with static or dynamics disorder. Differentiating among different models presents a challenge, as many distinct physical scenarios lead to similar experimentally observable behaviors such as anomalous diffusion and nonexponential relaxation. Here we show that information-theory-based analysis of single-molecule time series, inspired by Shannon’s work studying the information content of printed English, can differentiate between Markov (memoryless) and non-Markov single-molecule signals and between static and dynamic disorder. In particular, non-Markov time series are more predictable and thus can be compressed and transmitted within shorter messages (i.e. have a lower entropy rate) than appropriately constructed Markov approximations, and we demonstrate that in practice the LZMA compression algorithm reliably differentiates between these entropy rates across several simulated dynamical models.

## 1. Introduction

Single-molecule studies that track molecular conformations in real time have opened a new window on biomolecular folding, function of molecular machines, and other cellular phenomena. A critical limitation of such experiments, however, is that they usually track only a few quantities of interest over time, even though the observed behavior is driven by underlying physical processes described by orders of magnitude more degrees of freedom. In other words, the experimental data describes a low-dimensional projection of high-dimensional molecular dynamics. It is known that such projected dynamics are highly complex and often intractable, their common property being that they exhibit memory effects^1^; that is, they are non-Markov processes. Yet to interpret single-molecule data, phenomenological Markovian models are commonly invoked for observed quantities of interest *x*(*t*), such as diffusion^2, 3^ or evolution of *x* along a biased random walk on a lattice when *x* is discrete^4^. Signatures of non-Markovian dynamics such as anomalous diffusion, where the mean-square displacement of *x* grows nonlinearly with time^5, 6^, or nonexponential relaxation have been previously reported (see, e.g., refs.^7-13^), but the challenge, then, is to choose the right dynamical model out of the multitude of possibilities^5^. Data-driven Bayesian inference models of single-molecule time series have enjoyed considerable success in recent years^14-19^, but they usually require physical insight in order to constrain the space of possible models, and they, too, often assume that the observed dynamics is a one-dimensional random walk even if the number of discrete states is not specified a priori.

Is it possible to tell whether the observed signal can be explained by a Markov process, or whether a non-Markov model, or a model of a higher dimensionality, is called for by the data? For the case where the experimental observable *x* is a continuous variable, one such Markovianity criterion has been established recently^20^ based on analyzing spatial intervals [*a, b*] and comparing to theoretical inequalities the frequency with which the trajectory *x*(*t*) transitions through or loops back when entering the interval, but this approach is inapplicable to processes where the observed experimental quantity takes on discrete values. Other non-Markovianity signatures have been established^21, 22^, and an information-theoretic method based on computing the mutual information of the true dynamics and its Markovian model has been proposed^23^. For discrete-state processes, a solution was already described by Shannon in his classic work^24^ where he estimated the information content (i.e. the Shannon entropy) of printed English. Specifically, a printed text may be viewed as a discrete-time signal … *i*(*t* − 1), *i*(*t*), *i*(*t* + 1) …, where the discrete variable *i* ∈ {1, …, *N*} encodes the text characters (*N* being the alphabet size). In the simplest – and clearly unrealistic – language model, the text is a sequence of statistically independent characters, and its statistical properties are completely specified by the frequencies *P*(*i*), 1 ≤ *i* ≤ *N*, with which each character appears in the text. A binary representation of the alphabet requires log_2_ *N* bits per character, but by encoding frequently occurring characters with fewer bits and rare ones with more bits, one can compress the text, with the theoretical compression limit being given by Shannon’s entropy *H* (in bits per character):

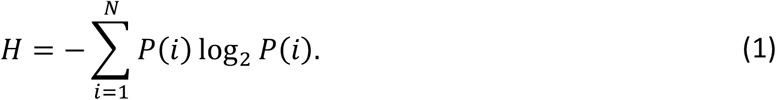

A better language model, however, recognizes that certain adjacent pairs of letters are more frequent than others. For example, “t” is more likely to be followed by “h” but not by “w”. This observation can be used to compress the text further, by encoding the text as a sequence of character pairs rather than individual letters. To quantify the benefit of this pair encoding, let *T*(*i* → *j*) be the conditional probability that the *i*-th character is followed by the *j*-th character. Then the string of letters produced by this model is a discrete-time (1^st^ order) Markov process, with *T* the transition probabilities. The theoretical limit for how much the text described by this model can be compressed is given by the *first-order entropy rate h*^(1)^,

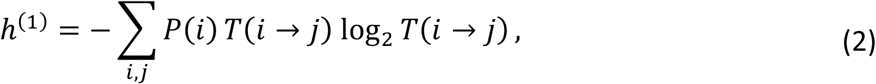

measuring the conditional entropy of knowing the next symbol given the previous one, and it can be easily verified that *h*^(1)^ ≤ *H*. (The entropy rate here is information-theoretic and is not to be confused with the thermodynamic entropy production rate, which quantifies a non-equilibrium system’s heat exchange with its environment.) Shannon then proceeded to generalize this idea to higher-order correlations by computing the conditional probability that each character appears given the two previous characters (a 2^nd^ order Markov process with entropy rate *h*^(2)^), and so on. The limit 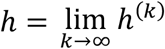 estimates the true information content (in bits per character) of the language.

A key observation is that *h*^(*k+1*)^ ≤ *h*^(*k*)^: a higher-order estimate of the entropy rate always yields a lower value than a lower-order estimate. Knowing more preceding characters can only help us to guess the next character more accurately, and thus the information revealed by the next character is lower. For similar reasons, the entropy rate of a non-Markov process is always bounded above by the entropy rate *h*^(*k*)^ of the *k*-th order Markov approximation of that process.

Although Shannon accomplished the remarkable feat of estimating the entropy of the English language without computers, we now can automate the same task by employing lossless compression algorithms. Given a sample path of a stationary stochastic process, the size (in bits per step) of the compression algorithm output is known to converge, asymptotically in the limit of large path size, to the theoretical entropy rate *h* of the process^25^. So, for example, to detect whether a piece of text – or a single-molecule trajectory – can be explained by first-order Markovian dynamics, we can on the one hand compress the text and compute the implied entropy rate *h* of the process from the compressed size; and on the other hand, measure the character-pair transition probabilities *T*(*i* → *j*) in the text and compute the first-order entropy rate *h*^(1)^ using Eq. 2. In principle, *h* ≈ *h*^(1)^ precisely when the original process is first-order Markov. In practice, of course, we need to worry that the compression algorithm is imperfect: it will overestimate *h*, and if the error is sufficiently large, the above procedure may fail to differentiate between Markov and non-Markov processes.

In what follows, we explore whether this “compression test” can reveal memory effects when applied to several families of non-Markov models commonly used to describe single-molecule dynamics. We find that this method can indeed differentiate between Markov and non-Markov dynamics. We further show that it can differentiate between static and dynamic disorder and thus narrow down the choice of a model to describe anomalous-diffusion-type phenomena. The rest of the paper is organized as follows: in Section 2 we describe the technical details of the method. Section 3 reports on its applications to several stochastic models. Section 4 concludes with further remarks and a discussion of future applications.

## 2. Methods

### 2.1. Core algorithm

We assume that the experimental time-dependent observable *i*(*t*) takes *N* possible discrete states, and that the process *i*(*t*) is stationary. We then present a general algorithm, based on the compression test outlined above, to determine whether the process is likely Markov or non-Markov. Our method can further distinguish between *k*-th order Markov processes, for modest *k*, and non-Markov processes with longer-term memory effects. We assume for now a discrete-time process and extend our approach to determining the Markovianity of continuous-time processes in section 2.2.

First, we estimate the entropy rate of the process. We compute a sample path *i*(1), *i*(2), …, *i*(*L*) of length *L* and represent it as an *L*-character string with an alphabet of size *N*. We compress the string using LZMA2 lossless encoding, as implemented in the xz library (https://tukaani.org/xz/). The size of the resulting compressed data is measured in bits and divided by *L* to obtain an estimate 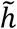 of the entropy rate. In our experiments we chose *L* ≈ 10^8^; see the supplemental material for full methodological details.

The entropy rate estimate is affected by compressor error: it will differ from the process’s “true” entropy rate *h* by an error 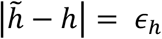 depending on the implementation details of the compression algorithm (which stores metadata within the output file) and the inability of the algorithm to perfectly compress the sample data. Although *ϵ*_*h*_ is guaranteed to converge to zero in the limit of an infinitely long sample path *L* → ∞, in practice the error remains significant even for paths containing on the order of a billion steps.

Next, we construct an approximate *k*-th order Markovian description of the process *i*(*t*). For any sequence of *k* consecutive states *S* = *i*(*m*), *i*(*m* + 1), …, *i*(*m* + *k* − 1) encountered in the sampled path and for any *i* we can estimate the (conditional) transition probability *T*(*S* → *i*) that *S* is followed by *i* from the frequency of the subsequence *S, i* within the sampled string. We then sample a path *j*(1), *j*(2), …, *j*(*L*) from the *k*-th order Markov process defined by these probabilities and compress it as described above, yielding entropy rate estimate 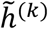. Note that this estimate does not exactly match the true entropy rate of the *k*-th order Markov process,

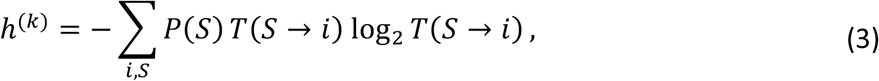

since again the estimate suffers from compressor error 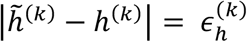. By comparing 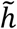 to 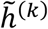, rather than to *h*^(*k*)^ directly, we exploit favorable cancellation of error: *if* the original random process is indeed *k*-th order Markov, then *ϵ*_*h*_and 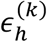 are drawn from the same distribution, so that 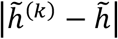 is small even for values of *L* for which the individual errors remain large.

Markovianity detection therefore amounts to checking if 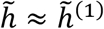. When 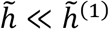 the process is likely non-Markov; higher-order Markov processes with longer memory can be discerned by repeating the experiment for different *k* (though the cost of doing so grows exponentially with *k*). We show below that this procedure does indeed clearly distinguish between Markov, second-order Markov, and non-Markov processes.

### 2.2. Generalization to continuous-time processes

The core algorithm can also be applied to continuous-time processes. We use kinetic Monte Carlo simulations^26-28^ to obtain stochastic trajectories *i*(*t*) reporting on the location of the system in the discrete state space as a function of time *t*. Depending on the problem (and as described below for each individual case in the next Section) we then either sample the continuous-time trajectory at finite (and sufficiently small) time intervals *Δt*, which results in discrete time series *i*(0), *i*(*Δt*), *i*(2*Δt*) … approximating the original trajectory, or describe the trajectory as a succession of states *i*_0_, *i*_1_, *i*_2_, …, where *i*_*k*_ is the first site visited by the continuous-time process after leaving the previous site *i*_*k*−1_. We will refer to the former description as “continuous-time” (even though *i* is sampled at finite time intervals) and the latter as “discrete time.” By construction, self-transitions *i* → *i* are not allowed in discrete-time processes, since each new state visited by the walk is different from the previous one. In contrast, the continuous-time description allows for self-transitions, where the state *i* remains the same during the sampling interval *Δt*, i.e., *i*(*t* + *Δt*) = *i*(*t*). It is of course possible that the process visits a state other than *i* and returns to the original state, resulting in *i*(*t* + *Δt*) being equal to *i*, though the probability of such events can be made arbitrarily small by choosing a sufficiently small value of *Δt*.

An important family of continuous-time signals consists of processes *i*(*t*) whose discrete-time description is Markovian but continuous-time description is not. Such processes are known as semi-Markov or renewal processes^29^, and they will be further discussed in Section 3.7.

## 3. Results

### 3.1. Single-file diffusion, Simple Case

A classic example of a random walk with (generally) long memory is single-file diffusion^30, 31^ (Fig 1a). This model has applications, for example, as the prototype of diffusion in crowded environment of a biological cell^5, 32^, passage of multiple solute particles across a biological channel^33^, and non-Markovian barrier crossing kinetics^34^. Here we use a discrete-time lattice formulation, in which *M* particles occupy discrete positions on a ring with *R* sites. A particle can move to an adjacent site if it is unoccupied, and each step of the single-file diffusion process consists of one such move (chosen uniformly at random). An observer monitors the location *i*(*t*) on the ring of a single tagged particle (shown in red in Fig. 1a), where the discrete time *t* enumerates successive steps of single-file diffusion. It is known that the motion of the tagged particle generally exhibits anomalous diffusion and thus *i*(*t*) is not a Markov process.

**Figure 1:**
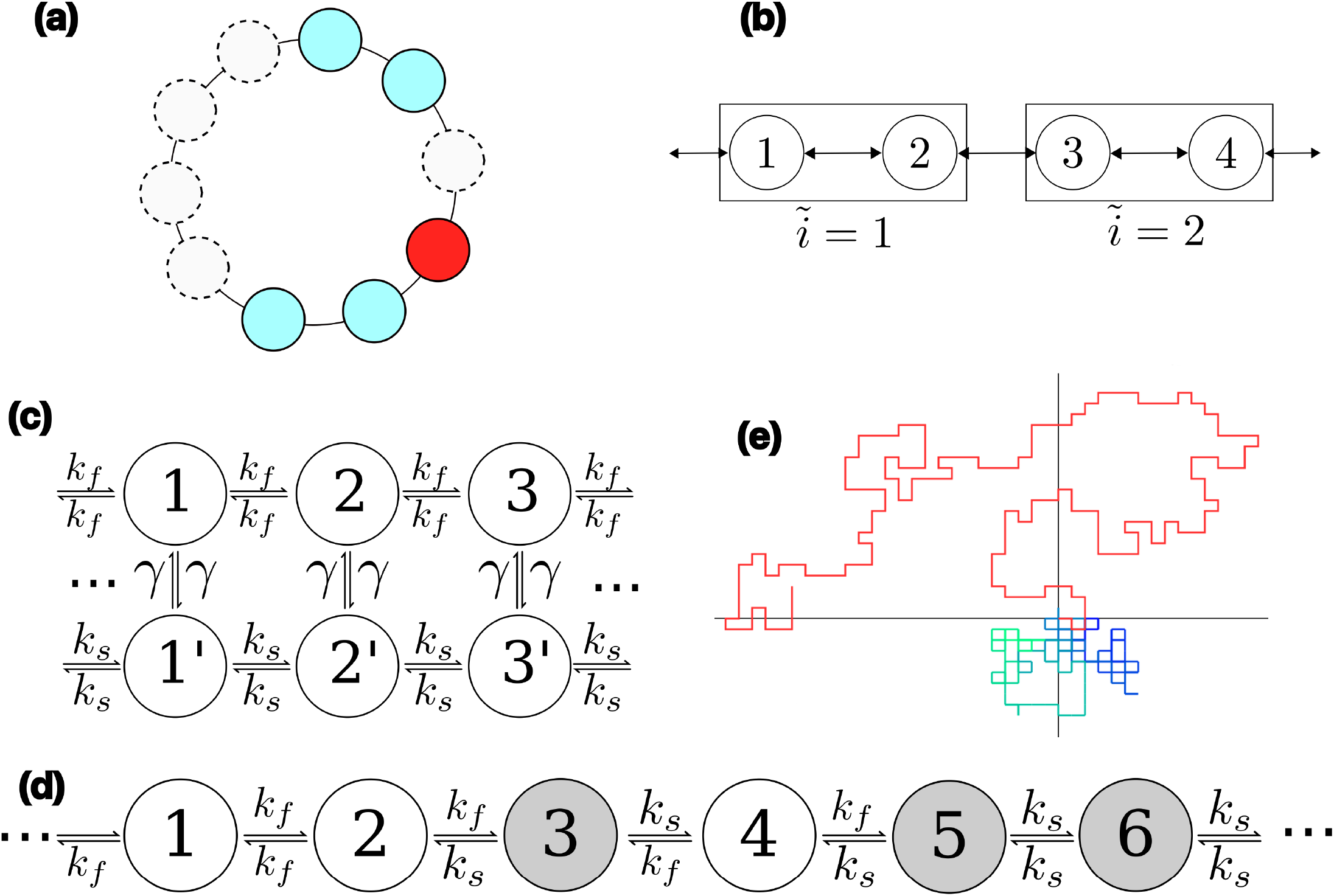
The models under study in this paper. (a) Single-file diffusion on a ring lattice. Each particle (filled circle) can move to an adjacent lattice site only when the latter is vacant (empty circle). The observer monitors the position of a single tagged particle (red). Here the number of lattice sites is *R* = 10 and the number of walkers is *M* = 5. (b) Coarse-grained random walk, in which every *n* adjacent lattice sites (n=2 here), are grouped into a single state, is a non-Markov process. The labels inside the circles enumerate lattice sites *i*, while the labels outside the boxes enumerate the coarse grained sates 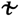. The observer can only tell which boxed state *ĩ* the particle is in. (c) A random walker with two internal states. The transition rates between pairs of states (1,2), (2,3), etc. have a higher value *k*_*f*_ than those between pairs (1’,2’), (2’,3’), *k*_*s*_, and the walker can switch between internal states (i.e. between states *n* and *n*′) with a rate *γ*. Thus the random walk is fast along the track formed by states 1,2,3,…, and slow along 1’,2’,3’….. The observer, however, cannot differentiate 1 from 1’, 2 from 2’, etc., and so the resulting observed process is non-Markovian. (d) A static-disorder model. Each site is randomly chosen to be fast or slow. Transitions leaving slow sites (gray) occur with rate *k*_*s*_, while transitions leaving fast sites (white) occur with rate *k*_*f*_ (e) Self-avoiding random walk on a 2D lattice. Shown is a trace of a self-avoiding walk (red) and a non-self-avoiding walk (blue-green) on a square lattice.

It is instructive to consider the simple, analytically tractable, case with *R* = 3 and *M* = 2 illustrated in Figure 2. The entropy rate *h*^(1)^ for the tagged particle using the 1^st^ order Markov process approximation can be calculated analytically: at any step, one of the two walkers moves, and thus the probability that the tagged particle moves is ½. When it does, it moves either clockwise or counterclockwise with the same probability, and thus, at any step we have 3 outcomes: no move 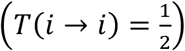, move clockwise 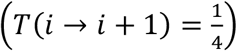, and move counterclockwise 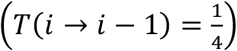. Using Eq. 2, the entropy rate based on these transition probabilities is 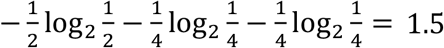 bits/step.

**Figure 2.**
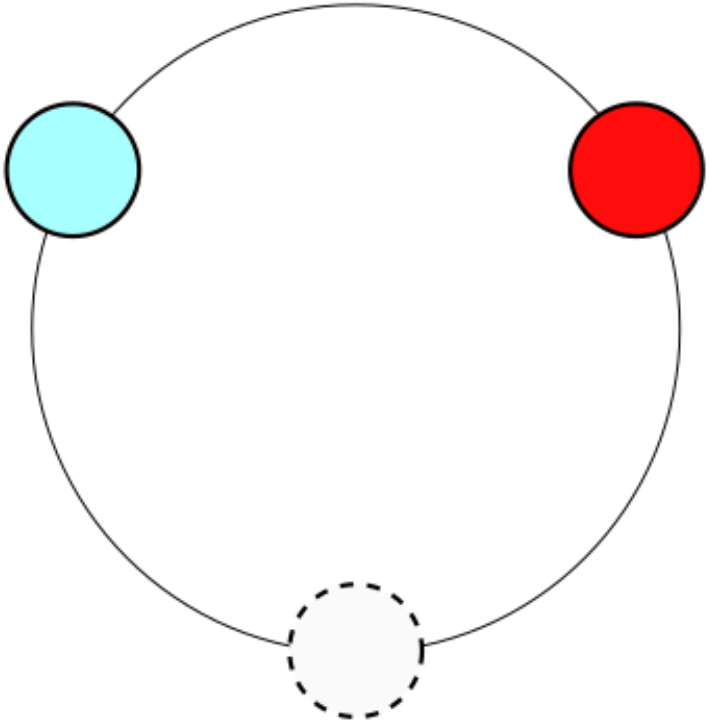
Single-file diffusion with *M* = 2 walkers and *R* = 3 sites on a ring, where one tagged walker (red) is being observed. The filled circles represent occupied sites, and the open circle depicts the vacancy (i.e. unoccupied site). At every timestep, a random walker is chosen to move, and moves into the vacancy, leaving its old site vacant.

The entropy rate *h*^(2)^ of the 2^nd^ order Markov process approximation can also be found analytically via a change of variable. Specifically, consider the process *ℓ*(*t*) = *i*(*t*) − *i*(*t* − 1). This process may be viewed as a “step” representation of the particle’s trajectory, as it informs about the direction and length of the step taken. Since there is a one-to-one correspondence between *i*(*t*) and *ℓ*(*t*) assuming that the starting point *i*(0) is known, the two processes have the same entropy rate. Suppose now that the tagged particle takes a step in the clockwise direction, *ℓ* = +1. It easy to see (Fig. 2) that the next step in the same direction would be into a site that is occupied, so *T*_*ℓ*_(+1 → +1) = 0, where we have written *T*_*ℓ*_ to emphasize that these are the transition probabilities of *ℓ*(*t*) instead of *i*(*t*). The tagged and untagged particles have the same probability to move, and so we find 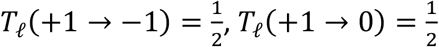. By symmetry, we also find *T*_*ℓ*_(−1 → −1) = 0, 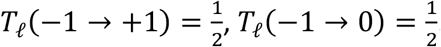. Using similar arguments, one can show that 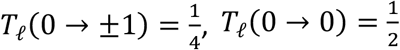. The unconditional step probabilities are evaluated in a similar manner, 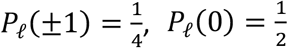. Using Eq. 3 we now find 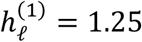 bits/step. Again, the subscript *ℓ* indicates that we are considering the process *ℓ*(*t*) rather than *i*(*t*). Since *i*(*t*) is recoverable from *ℓ*(*t*), and *ℓ*(*t*) can only depend on *i*(*t*) and *i*(*t* − 1), 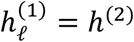, and hence *h*^(2)^ = 1.25 bits/step.

Note that we have derived the 2^nd^-order Markov entropy rate of the process *i*(*t*) by computing the 1^st^-order entropy rate of a different representation *ℓ*(*t*) of the same physical process. This highlights the fact that different representations of the same system may have different Markov orders. In our examples below that take place on a structured lattice, we will continue to make use of the “step” representation *ℓ*(*t*) = *i*(*t*) − *i*(*t* − 1) when convenient; the entropy rates for the two representations are related by 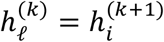. We will often simplify notation and use 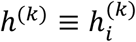 to denote the entropy rate of the process in the original “lattice site” representation (in accord with the usual convention in physics that a simple random walk depending only on the current site is regarded as a 1^st^ order Markov process), but will use a subscript (e.g. *i* or *ℓ*) when we want to emphasize which representation is used to calculate the order.

Although *h*^(2)^ provides an improved estimate of the single-file diffusion entropy rate, it is still an overestimate. The true entropy rate is *h* = 1 bit/step, as can be seen from the following argument: there is a one-to-one correspondence between sample paths *i*(*t*) of the tagged particle, and sample paths *v*(*t*) of the vacant site (dashed circle in Fig. 2). But the latter is a Markovian, unbiased random walk (since at every step it exchanges places with one of its two occupied neighbors), and so its entropy rate is exactly *h* = 1 bit/step. Note that this example again shows how the same random process can be Markov in one representation (motion of the vacancy) and non-Markov in a different one (motion of the tagged particle).

Applying compression to a trace with *L* = 10^9^ steps gives the estimates 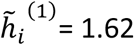 bits/step, 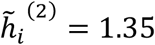 bits/step, and 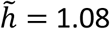 bits/step. The discrepancy between *h* and 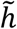 stems from the compressor error 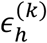 vanishing very slowly as a function of *L*. If we subtract a constant *Δh* ≈ 0.10 from all three measurements, we find that 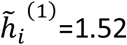 bits/step, 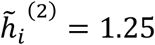 bits/step, and 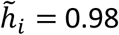 bits/step, which is almost perfect agreement with the theoretical values.

### 3.2 Single-File Diffusion, General Case

For larger *M* and *R*, we use the lattice-site representation to obtain compression-based entropy-rate estimates 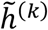 for several values of *k*, and compare them to the compression-based estimate 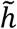 of the true entropy rate, as well as to the entropy rates *h*^*(k)*^ (Eqs. 2-3) of the Markov approximations in the absence of any compressor error.

Results are shown in Figure 3a for *M* = 7 walkers and for different numbers of sites *R*. Here, as in the *M* = 2, *R* = 3 case, we observe a discrepancy between *h*^*(k)*^ and 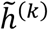. As discussed in Section 2, it is most meaningful to compare the 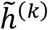 against each other and against 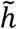.

**Figure 3.**
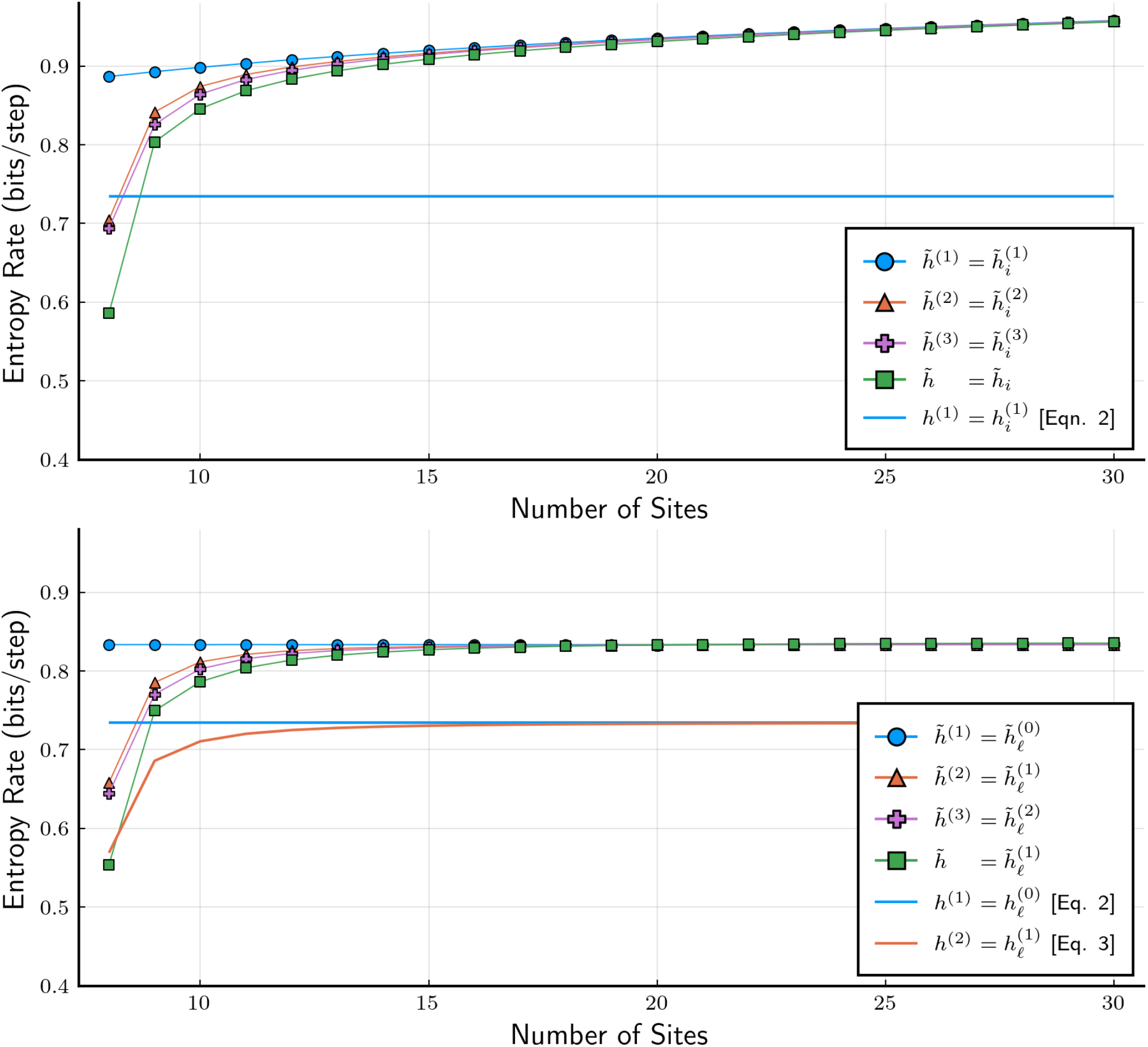
Entropy rates of a tracer particle in a single-file diffusion setup on a ring lattice shown as a function of the number *R* of lattice sites for a fixed number of random walkers, *M* = 7. Quantities with a tilde (e.g. 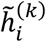) were computed using compression, while quantities without a tilde (e.g. 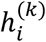) were obtained using Eqs 2 or 3 using numerically estimated transition probabilities. When the number of open sites *R* − *M* is low, the system behaves in a very non-Markov manner. Once the number of sites greatly exceeds the number of particles, the system’s behavior approaches Markov.(a) Entropy rates computed using the site-based *i*(*t*) representation. (b) Entropy rates computed using the step *ℓ*(*t*) representation.

When *M* ≈ *R*, we further observe that the true entropy rate 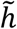 is significantly lower than its finite-order estimates 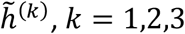, indicating strongly non-Markovian character of single-file diffusion. Since 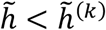, the memory of the previous *k* states is insufficient to construct an adequate description of the process. As *L* increases, however, the “clashes” between walkers become increasingly unlikely, and each walker diffuses freely in the limit *R* ≫ *M*, thus undergoing Markovian dynamics. Consistent with this observation, the true entropy rate estimate 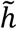 and its *k*-order Markovian estimates 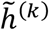 converge to the same value as *R* increases. We also note that, as expected, the estimated entropy rates decrease with increasing order *k*, that is, 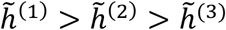.

We now compare the entropy-rate estimates based on the site representation *i*(*t*) and on the step representation *ℓ*(*t*). In theory, both processes should have the same entropy rate *h*, and, moreover, we have 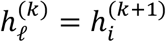. The compression-based estimates of these quantities, which are greater than their true values, are however not guaranteed to be the same, and indeed they are not (compare Fig. 3b with Fig. 3a). Moreover, compression-derived entropy rates 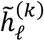 are significantly greater than their counterparts 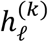 obtained from Eqs. 2-3 using numerically estimated transition probabilities (Fig. 3b). Importantly, however, the *relative* orderings between 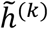 are preserved. Thus examination of either Figure 3a and Figure 3b unambiguously leads to the representation-independent conclusions (1) that single-file diffusion is a process with long memory that cannot be described by *k*-order Markov process with *k* ≤ 3 when most of the lattice sites are occupied (*M* ≈ *R*) and (2) that the process becomes increasingly close to a Markov process as *R* grows much larger than *M*.

### 3.3. Memory in coarse-grained random walks

A fundamental source of dynamical memory is coarse-graining^1^. An experiment cannot resolve each of the individual microscopic states of the molecule, so multiple microscopic states are effectively lumped into collective observable states. To illustrate this effect, we use a simple toy model in which the true dynamics are an unbiased discrete-time random walk along a one-dimensional lattice, whose sites are enumerated by *i*. Suppose that the spatial resolution of the experiment only allows one to resolve a length of *n* adjacent sites. The observations thus form a random process on groupings of *n* adjacent sites (Fig. 1b). For example, sites *i* = 1,2, … *n* correspond to a collective state *ĩ* = 1, sites *i* = *n* + 1, *n* + 2, … 2*n* to *ĩ* = 2, etc. The new variable *ĩ* also undergoes an unbiased (in the sense that the unconditional probabilities of making steps left and right are the same) discrete-time random walk, but this walk is no longer Markovian: as will be seen below, the conditional probability of making a step left or right depends on the direction of the preceding step.

We cannot directly apply our method to compute 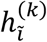, as we did in Section 3.2, since (as stated in Section 2) our method requires a random process on finitely-many states. However, we can once again define *ℓ*(*t*) = *ĩ*(*t*) − *ĩ*(*t* − 1), which has the same entropy rate as *ĩ*(*t*), and examine 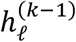 to determine 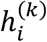. Given that the probabilities of stepping left *ĩ* → *ĩ* − 1 or right *ĩ* → *ĩ* + 1 are the same, we have *P*_*ℓ*_(−1) = *P*_*ℓ*_(+1) = 1/2, the same as for the original random walk. Therefore, the first-order Markov approximation of *ℓ*(*t*) has entropy rate of *h*^(1)^ = log_2_ 2 = 1 bit per step (Eq. 2). Intuitively, at every step one needs to specify whether the walker moves left or right, which occur with equal probability. The coarse-grained walk, however, is not Markovian because the direction of each step is correlated with the direction of the previous step. For example, for *n* = 2, one can show that the conditional probability of making the next step in the same direction as the previous step is 1/3 while the probability of making a step in the opposite direction is 2/3. In other words, we have

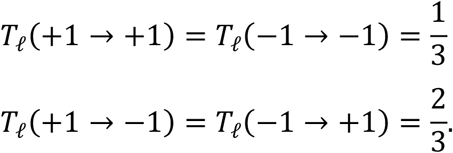

Moreover, it is easy to see that the direction of a step only depends on that of the previous step and not on earlier steps. In other words, the 1^st^ -order Markov model in the step representation (2^nd^-order in the site representation) is exact for our coarse-grained random walk.

Using Eq. 2 and the fact that the (unconditional) probabilities to make a step in either direction are *P*_*ℓ*_(±1) = 1/2, we find an entropy rate 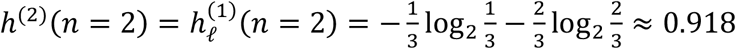 bits/step. In contrast, when we applied the compression method to a synthetic 2^nd^ order Markov walk generated according to the above transition probabilities (using *L* = 10^8^ steps) we obtained a value of 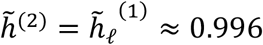 bits/step. Once again, the compressor overestimates the true entropy rate— in fact, applying the compressor to the original, ungrouped random walk (which has an entropy of exactly 1 bit/step) yields a result of 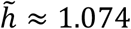 bits per step, a value of *ϵ*_*h*_ = 0.074 bits/step higher than expected. However, 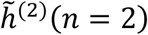 and 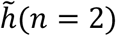 show excellent agreement, and this comparison reveals that the grouped random walk is second-order Markov (see Fig. 4a).

**Figure 4.**
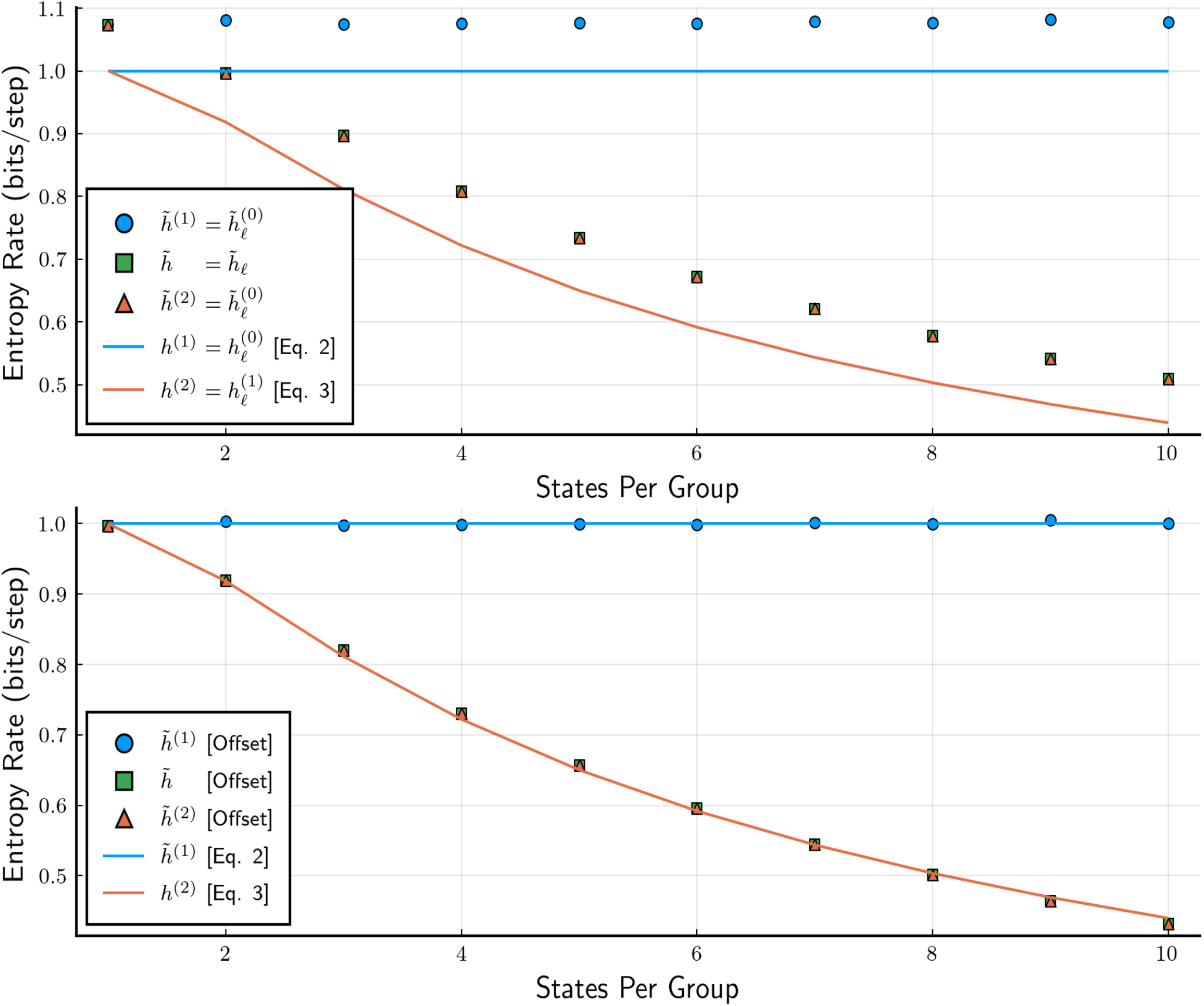
(a) Entropy rate (per step) 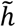 of a coarse-grained random walk as a function of the number of states *n* forming a single coarse-grained state. This entropy rate is compared with the compression-derived entropy rates 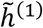 and 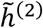 of the 1^st^ and 2^nd^ order approximating Markov processes, as well as with their theoretical values *h*^(1)^ = 1 bits/step and *h*^(2)^ given by Eq. 4. (b) The same plots after subtracting the offset *Δh* from the compressor-derived entropy rates.

More generally, for coarse states containing *n* adjacent sites, one can show that

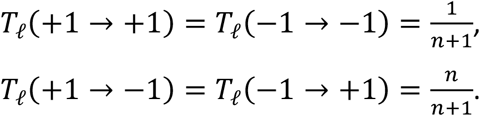

Intuitively, a walker entering a group of *n* states from the left is much more likely to escape it back to the left than to cross this group and escape to the right. As a result, the entropy rate of the second-order Markov approximation is

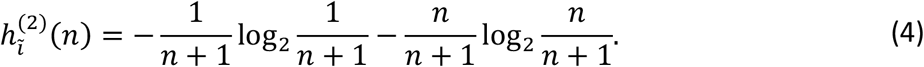

Figure 4 compares the compression-derived entropy rate 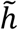 with the Markovian approximations calculated in two ways: (1) using the exact theoretical values known in this case (i.e., *h*^(1)^(*n*) = 1 bit/step and *h*^(2)^(*n*) given by Eq. 4 and (2) using our compression method. As with the single-file diffusion system, the compression algorithm does not attain the theoretical compression ratio and overestimates the entropy rate in both cases, but comparison of 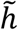 and 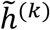 reveals that the system is second-order Markov. Moreover, one observes from Figure 4a that 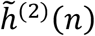 differs from *h*^(2)^(*n*) by a nearly constant offset. Subtracting this offset from 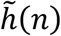 results in an estimate for the entropy of the coarse-grained random walk that is virtually indistinguishable from the true value (Fig. 4b).

### 3.4. Random walker with internal states

Another example of coarse-graining is the model shown in Fig. 1c. Unlike the previous case, we are now considering a continuous-time process, where a random walker can be in one of the two internal states; in one, its diffusion is fast (quantified by a jump rate *k*_*f*_) and in the other it is slow (jump rate *k*_*s*_). The walker can switch between the two internal states stochastically, with a switching rate *γ*. The kinetic scheme (Fig. 1c) thus consists of a “fast” and a “slow” track, with the walker randomly switching between the two. According to this scheme, the time evolution of the probabilities *P*(*i, t*) and *P*(*i*′, *t*) to occupy sites *i* and *i*′ obey the continuous-time master equation,

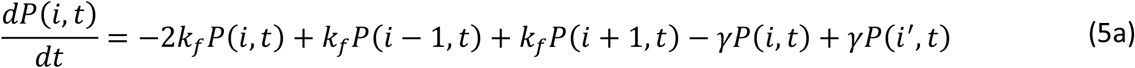

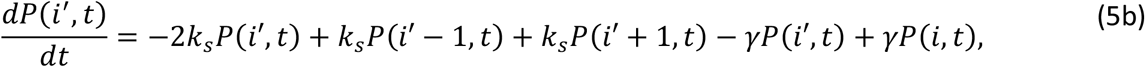

which describes a Markov process. The states *i* and *i*′, however, appear indistinguishable to the observer and are thus lumped to a single coarse state characterized by the apparent position *i* = 1,2, … of the walker along the track. Unless *k*_*f*_ = *k*_*s*_, the time evolution of the probability *P*(*i, t*) + *P*(*i*′, *t*) to be at position *i* cannot be described by a master equation like Eq. 5a or 5b: the observed random walk is non-Markovian.

Models of this type have been used, e.g., to describe the dynamics of biomolecular motors traveling along their tracks^4, 35, 36^, even though in our case the walker has no directional bias and thus does not undergo unidirectional motion characteristic of molecular motors.

The stochastic time evolution corresponding to Eq. 5 was simulated using the kinetic Monte Carlo method (see, e.g., refs.^26-28^) and the entropy rate was estimated using the “continuous time” approach described in Section 2.2, using a time step *Δt* chosen to satisfy the inequalities *k*_*f*_*Δt* ≪ 1, *γΔt* ≪ 1, to guarantee that the probability of the walker changing state more than once during *Δt* is negligibly small. As with the coarse-grained random walk, we computed entropy rate using the step representation (and make use of 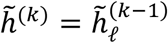) to avoid dealing with infinite sites.

The compression-derived entropy rate of this walk is shown in Figure 5a as a function of the switching rate *γ* for the case *k*_*s*_ = 0.1*k*_*f*_, and compared to the entropy rates of the first- and second-order Markov approximations. For all values of the switching rate *γ*, the true entropy rate of the walker is always lower than that of the two reference Markov processes (whose entropy rates are indistinguishable), indicating non-Markovianity of the underlying process. Despite the significant statistical noise observed in Fig. 5a, particularly at slow switching rates, the difference between the true entropy rate and that of the first-order reference Markov process (shown in Fig. 5b) is always negative and is less noisy than the absolute entropy rates shown in Fig. 5a – thus the compression method detects the non-Markovianity of the underlying dynamics reliably even when the simulations have not fully converged.

**Figure 5.**
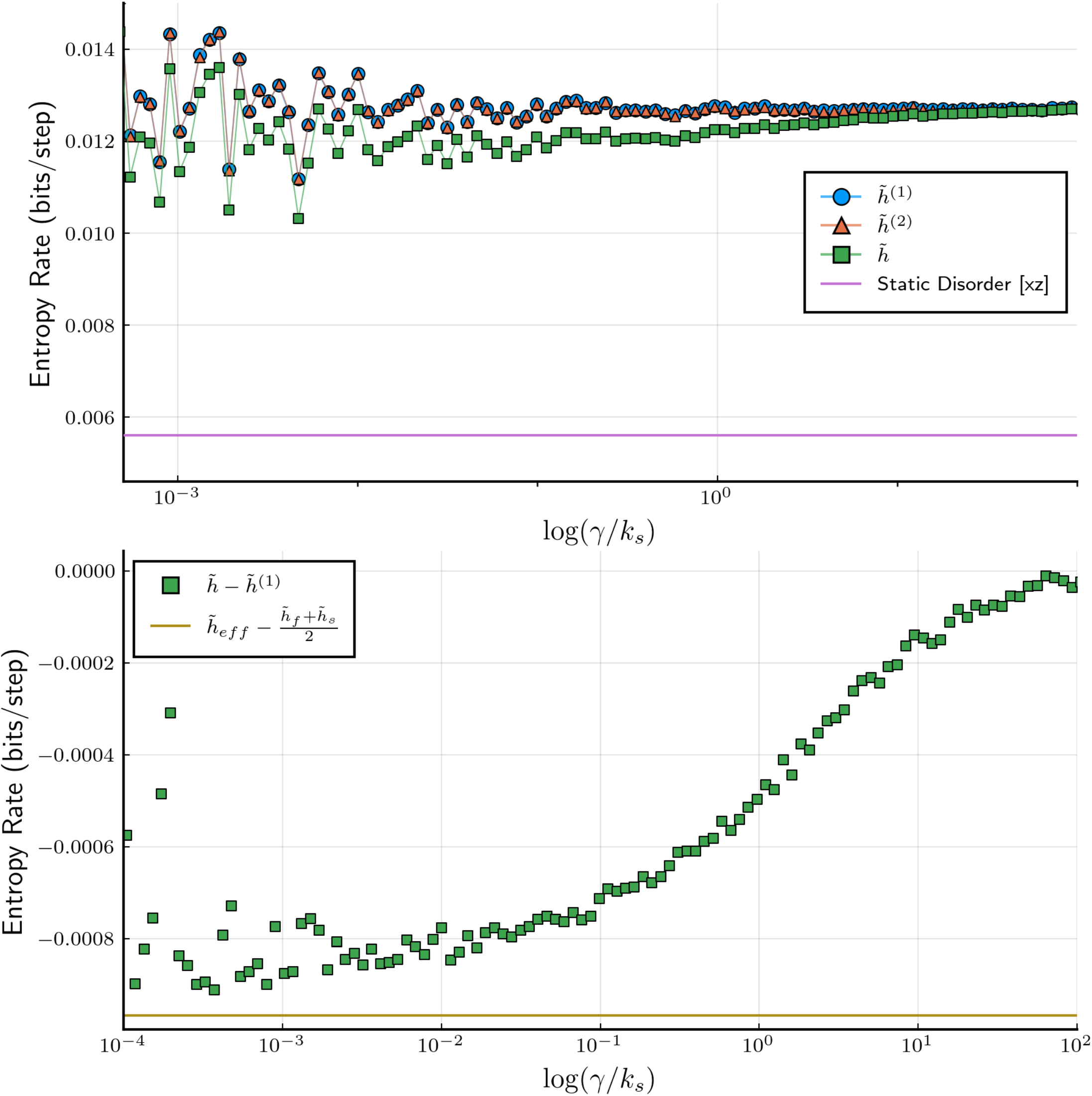
(a) Entropy rate for the random walk with two internal states (Fig. 1c), plotted as a function of the switching rate *γ* between the fast and slow tracks and compared with the reference 1^st^ and 2^nd^ order Markov processes. Solid line represents the compression-derived entropy rate for the random walk with static disorder (Fig. 1d). The kinetic parameters are *k*_*s*_ = 0.1, *k*_*f*_ = 1.0. (b) The difference *h* − *h*^(1)^ between the compression-derived true entropy rate and that of the Markovian approximation plotted as a function of the switching rate. At low switching rates, *γ* → 0, the entropy rate (measured relative to its *γ* → ∞ limit) is seen to approach the expected value equal to the mean of the (compression-derived) entropy rates of two Markovian processes, the slow one (with the jump rate *k*_*s*_) and the fast one (jump rate *k*_*f*_).

The dependence of the entropy rate on the switching rate also agrees with what one expects on physical grounds. Specifically, in the limit of high switching rates, *γ* ≫ *k*_*f*_, the process becomes effectively Markovian, with the effective transition rate coefficients between adjacent positions on the track equal to the average rate, *k*_eff_ = (*k*_*s*_ + *k*_*f*_)/2, with a corresponding entropy rate of *h*_eff_. Note this Markov process is identical to the 1^st^ order reference Markov process, which is the same regardless of the switching rate. Consistent with this observation, we observe that (1) the compression-derived value 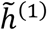 is independent of the switching rate (aside from statistical noise) and (2) the true entropy rate approaches the entropy rate *h*^(1)^ of the reference Markov process at large values of *γ* (Fig. 5a,b).

Consider now the opposite case of slow switching rate, *γ* ≪ *k*_*s*_. In this case we expect that the trajectory of the random walker will consist of long segments where the walker stays on the fast (or slow) track, jumping between neighboring lattice sites with a rate of *k*_*f*_ (or *k*_*s*_). Since the fractions of time spent on each track are 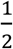, we expect the entropy rate to approach an average value of *h* = (*h*_*f*_ + *h*_*s*_)/2, where *h*_*f*_ (*h*_*s*_) is the entropy rate of a Markovian random walk along the fast (slow) track (i.e. a 1D random walk with a transition rate *k*_*f*_ (*k*_*s*_). Indeed, this is the behavior observed in Fig. 5b, with a compression-derived estimate of (*h*_*f*_ + *h*_*s*_)/2 agreeing with the compression-derived true entropy rate 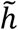 at low switching rates.

### 3.5. Compression differentiates between static and dynamic disorder

The two-track model considered in Section 3.4 is an example of a model with dynamical disorder, where the (mean) lifetime of the random walker on a lattice site can be either long (1/(2*k*_*s*_)) or short (1/(2*k*_*f*_)) depending on a dynamical variable (the internal state of the walker) that is itself undergoing a stochastic process switching between the slow and fast states. By construction, the probabilities of finding the walker in the slow and fast states are equal. It is instructive to consider a seemingly similar model with static (quenched) disorder. In this model of a 1D random walk (Fig. 1d), each lattice site is randomly assigned to have either long or short average dwell time. Transitions that leave the former occur with rate *k*_*s*_, and transitions that leave the latter occur with rate *k*_*f*_. Since in both models the random walker visits the “slow” and the “fast” sites with equal probabilities, one often uses these two types of models interchangeably, but the two models are not equivalent^37^. More generally, various anomalous-diffusion-type phenomena are often interpreted in terms of random walks on a network of “traps”, with a certain distribution of the trap energies leading to a distribution of trapping times^5^. One often models such a random walk as a renewal process (see Section 3.7 for further discussion of such processes), in which the times spent on each trap are statistically independent, but such an assumption is not correct when the trap properties are “frozen” in time^5, 37^.

While differentiating between static and dynamic disorder in anomalous diffusion phenomena is often a challenge, the compression-based method explored here achieves this goal rather straightforwardly, at least for the model system considered. Specifically, the entropy rate corresponding to the process with static disorder is always lower than that for dynamic disorder, as can be observed in Fig. 5a. It is easy to understand why: every time the random walker visits a new lattice site, the information gained consists of the site identity and of the time spent on this site. If the walker visits *the same* site again, less information will be gained in the case of static disorder, as information can already be inferred about this site’s dwell time from the time spent on this site in the previous visit.

### 3.6 Self-avoiding random walks

A random walk that is not allowed to cross its prior path offers an interesting example of a non-Markov process with infinite (or, more precisely, as long as the walk itself) memory. Such random walks have been studied extensively in the context of polymer physics, where they serve as models of polymer backbones. Moreover, a compression-based approach to the mathematically equivalent problem of computing the entropy of a polymer has been recently studied by Avinery, Kornreich and Beck^38^.

Here we consider self-avoiding 2D walks on a square lattice (Fig. 1e). Because of the symmetry between up/down and left/right directions, the probabilities of making step in one of the 4 possible directions are equal, *T*(*i* → *i*′) = 1/4, for any of the 4 sites *i*′ adjacent to *i*. Thus the 1^st^ order Markov model of the process is simply the random walk with the self-avoidance condition removed. For such a walk the number of possible step directions at each step is 4 = 2^2^, and thus its entropy rate is *h*^(1)^ = log_2_ 4 = 2 bits per step.

The 2^nd^ order Markov model of the self-avoiding walk clearly must account for the fact that any step cannot be the reverse of the previous step. Since there are only 3 possible steps left, a naïve estimate of the entropy rate would be *h*^(2)^ = log_2_ 3 ≈ 1.585 bits/step, assuming that all three possible directions are equally probable. A more precise estimate for this value, obtained numerically from Eq. 3 using transition probabilities measured from simulation data, is slightly lower, *h*^(2)^ ≈ 1.578. Note that this model is locally self-avoiding, since by demanding that a step cannot be reversed, we automatically ensure self-avoidance for 3 steps in a row. Thus, only long-range correlations (longer than 3 steps) may account for the difference between *h*^(2)^ for the 2^nd^ order Markov model and the entropy rate of the true self-avoiding walk. As we show below, the compression-based method is still capable of differentiating between these two models.

Figure 6 shows the entropy rates estimated numerically as functions of the walk length. To apply our method we sampled 250,000 length-*N* self-avoiding random walks using Sokal’s pivoting algorithm^39^. For sufficiently large values of *N*, the entropy rate becomes independent of the length of the walk. This property is an established fact^40^ but not immediately obvious, as the memory of the walk is as long as the length of the walk itself. Specifically, the total number of self-avoiding random walks asymptotically grows as

**Figure 6.**
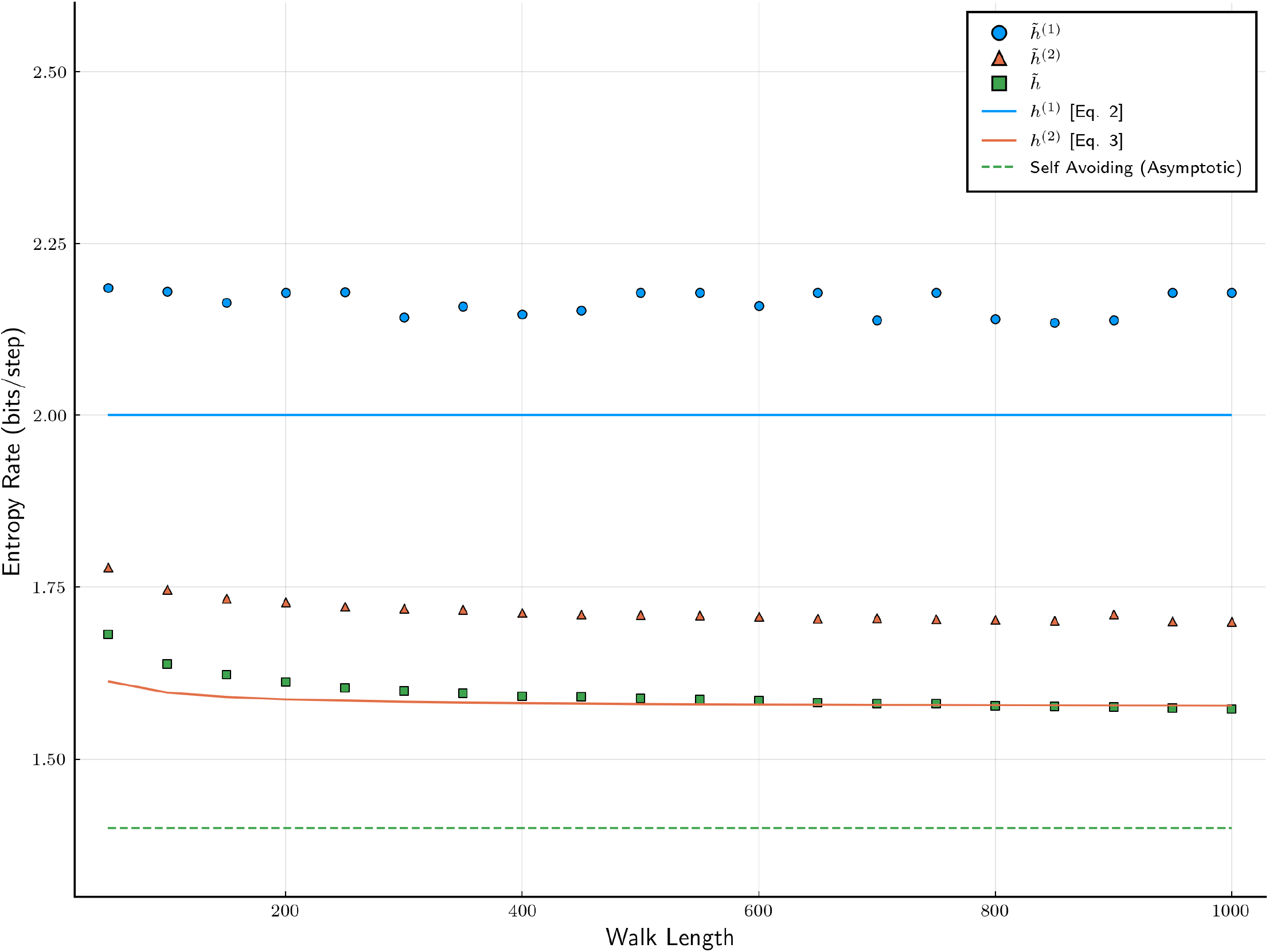
Entropy rates for self-avoiding walks of various lengths, compared with the with the entropy rates of the corresponding 1^st^ order and 2^nd^ order Markov models. Literature value for the entropy of the self-avoiding random walk^40^ is shown as a dashed line.

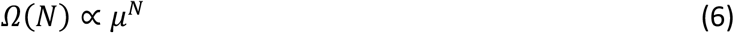

as *N* → ∞, where numerical estimates exist^41^ for lower and upper bounds for *μ*, 2.625622 < *μ* < 2.679193. We can interpret *μ* as the average number of possible step directions available at each step of the walk, and thus its entropy rate is

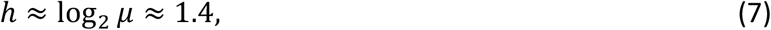

which we take as the “exact” entropy rate in this case. As seen in Figure 6, the compression method differentiates among the two reference Markov approximations and true dynamics, indicating that the actual observed process has more memory than that of one previous step (in fact of 3 previous steps given that the self-avoidance is guaranteed for any 3 consecutive steps in this model).

While the compression-derived estimate of the connectivity parameter, 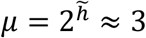, exceeds the best currently-known upper bound on the theoretical value, this estimate can be improved by accounting for the compression error: for the second-order Markov reference process we can compute 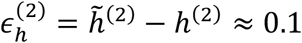 bits from Eq. 3 Assuming that 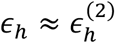 and applying this same correction to estimate 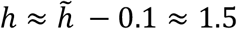 bits/step and *μ* ≈ 2.8, still somewhat greater than the known bound.

### 3.7. Continuous-time random walks (CTRW)

constitute an important class of non-Markov models used widely to describe anomalous-diffusion-type phenomena^6, 29, 42^. Milestoning, a method for computing long-time molecular dynamics, also uses a CTRW-type representation^43^. CTRW is an example of a semi-Markov or renewal process. A continuous-time random walker makes a step from a current site *i* to new sites *j* according to the conditional probabilities *T*(*i* → *j*), with the sojourn time at *i* before making this step drawn from a specified probability density *ρ*(*t*) (the case where this probability density is itself state dependent is a straightforward generalization). Thus the random walk can be viewed as a combination of two processes: the first one is the sequence of sites visited *i*_1_, *i*_2_, …, *i*_*n*_, … and the other is the sequence of sojourn times *t*_1_, *t*_2_, …, *t*_*n*_, … spent on each site. The first of the two is a discrete-time Markov process. The total entropy rate is the sum of the entropy rate for the discrete-time Markov process and the entropy rate associated with the sequence of the sojourn times. We will now focus on the latter. As the sojourn times are statistically independent, the entropy rate is just the entropy of the distribution *ρ*(*t*). More precisely, if we discretize the time by binning the temporal data into sufficiently fine time bins of equal size *Δt*, then the entropy rate is given by

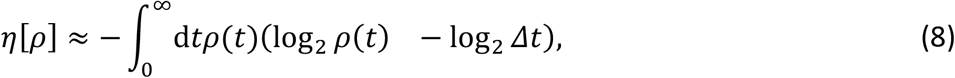

where the last term accounts for the dependence of the entropy on the resolution with which the time is measured.

Using the method of Lagrange multipliers, one finds that of all the distributions *ρ*(*t*) with a fixed first moment, 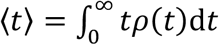, the one that maximizes *η*[*ρ*] is the exponential distribution,

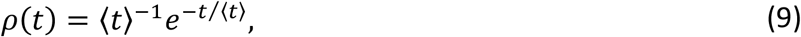

for which

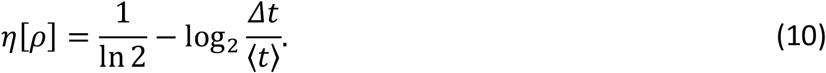

This distribution corresponds to a Poisson process, which is Markovian. This illustrates the general observation that Markovian processes maximize the entropy rate, provided that the average frequency of the transitions is fixed.

## 4. Discussion

Reconstruction of the underlying models of single-molecule dynamics from experimental observables has received much recent attention (see, e.g., ref.^44^ for a review) and remains a challenge in the field. The problem is further confounded by experimental constraints such as finite temporal and spatial resolution. In all of the examples studied here, compression-derived entropy rate estimates could differentiate between simulated Markov and non-Markov time series, as well as between models with dynamical and static disorder, even when the statistical errors or systematic errors introduced by the compression algorithm exceeded the difference between the entropy rate of the true signal and its candidate model (e.g. Markov description). We note that this method does not require a high data sampling rate that may be unattainable experimentally: indeed, the entropy rate at an arbitrary data sampling rate can be compared with that of the corresponding reference Markov process, the latter being always computable via Eqs. 2-3.

Markovian approximations to the non-Markov systems studied here provide upper bounds on the true entropy rate. In general, one can construct the *k*^th^-order Markov reference model of the process and to estimate its entropy rate *h*^*(k)*^; if *h*^*(k)*^ converges toward the entropy rate *h* of the experimental signal as *k* increases sufficiently fast, this result is a practical estimate of the number of previous steps needed to be included to produce an accurate description of the signal. For example, for the coarse-grained random walk with grouped states (Section 3.3) the 1^st^ order Markov process does not reproduce the “experimental” entropy rate, but the 2^nd^ order one (which happens to be the exact description in this case) does (Fig. 2). In contrast, the 2^nd^ order Markov model does not provide a noticeable improvement over the 1^st^ order one for the random walk from Section 3.6, suggesting long memory (as compared to the sampling time step). In such a case, a different description of the signal is desirable.

This work focused on signals where the observed variable is discrete. Such signals are common in single-molecule fluorescence studies in which the arrival of an individual photon can be viewed as a discrete event^45-48^. In contrast, force spectroscopy studies measure continuous variables such as a molecule’s extension or position of a molecular motor along its track^4, 49, 50^. Likewise, single-particle tracking studies measure continuous signals, see, e.g., ref.^51^. Can the compression method be applied to a continuous signal? A naïve answer to this is to digitize the observed variable by measuring it with a finite resolution. The resulting entropy rate is known as the “epsilon entropy” *h*(*ϵ*) (the parameter *ϵ* quantifying the resolution), which can be viewed as an approximation to the Kolmogorov-Sinai entropy^52, 53^. For stochastic signals, *h*(*ϵ*) is known to diverge as *ϵ* → 0, and its behavior for certain stochastic models has been studied^53^. To our knowledge, the practical utility in using *h*(*ϵ*) to differentiate between stochastic processes with and without memory has not yet been explored, and it will be the subject of our future work.

## Acknowledgments

We are grateful to Hagen Hofmann for many discussions. Financial support from the Robert A. Welch Foundation (Grant No. F-1514 to DEM), the National Science Foundation (Grant Nos. CHE 1955552 to DEM and IIS 1910274 to EV), and Adobe Inc. is gratefully acknowledged.

